# Shorter constant work rate cycling tests as proxies for longer tests in highly trained cyclists

**DOI:** 10.1101/2021.10.12.464126

**Authors:** Chantelle du Plessis, Mark Andrews, Lachlan J G Mitchell, Jodie Cochrane Wilkie, Trish King, Anthony J Blazevich

**Author notes:** **Corresponding author** Chantelle du Plessis, MSc.

## Abstract

Severe-intensity constant work rate (CWR) cycling tests are useful for monitoring training progression and adaptation as they impose significant physiological and psychological strain and thus simulate the high-intensity competition environment. However, fatiguing tests require substantial recovery and may disrupt athlete training or competition preparation. Therefore, the development of a brief, minimally fatiguing test providing comparable information is desirable.

**Purpose:** To determine whether physiological variables measured during, and functional decline in maximal power output immediately after, a 2-min CWR test can act as a proxy for 4-min test outcomes.

**Methods:** Physiological stress was monitored and pre-to-post-CWR changes in 10-s sprint power computed (to estimate performance fatigability) during 2- and 4-min CWR tests in high-level cyclists.

**Results:** The 2-min CWR test evoked a smaller decline in sprint mechanical power (32% vs. 47%, *p*<0.001), however both the physiological variables and sprint mechanical power were independently and strongly correlated between 2- and 4-min tests. Differences in V^·^O_2peak_ and blood lactate concentration in both CWR tests were strongly associated with the decline in sprint mechanical power.

**Conclusion:** Physiological variables measured during, and the loss in sprint mechanical power measured after, a severe-intensity 2-min CWR test were less than in the 4-min test. Yet strong correlations between 2- and 4-min test outcomes indicated that the 2-min test can be used as a proxy for the longer test. Because shorter tests are less strenuous, they should have less impact on training and competition preparation and may therefore be more practically applicable within the elite performance environment.

## Introduction

Regular monitoring of both athlete performance and overall fatigue levels are necessary to maximise physiological adaptation, manage recovery and training load, and to test the effects of interventions such as technique or equipment alterations. To do so, specific tests with sufficient precision are needed to detect relevant and meaningful changes. Appropriate testing protocols that can be used frequently without negatively impacting subsequent, prescribed training sessions or competition preparation are therefore required. Constant work rate (CWR) cycling tests are widely used to assess and monitor physiological and functional performance capacity in various athlete, clinical and rehabilitation environments. In athlete environments, these tests are typically performed at intensities within the severe domain (i.e., greater than the critical power threshold) for durations ranging 2-15 min (1). Their aim is to evoke a maximal effort that achieves the highest sustained average power output for the duration of the test, with minimal or no functional decline in mechanical power.

In cycling, an athlete’s ability to generate high power outputs during these tests should ultimately translate to a faster bicycle velocity in the field (e.g., during a time trial), which is critical to performance success. However, the maintenance of a high mechanical work rate (power output) requires a high muscle contractile force production at fast muscle shortening speeds, and therefore a high adenosine triphosphate (ATP) turnover rate and metabolic energy cost (2-4). The associated unstable metabolic environment can impact function at multiple sites within the muscle excitation-contraction coupling process at the sarcomere level and influence efferent output by the central nervous system (for reviews see (5-7)). Regardless of the mechanism, muscle function will be compromised irrespective of a participant’s voluntary effort capacity when exercising within the severe-intensity domain, leading to fatigue (8-11).

To monitor and maximise physiological adaptation, it is necessary to understand the effects of the fatiguing CWR test on the energetic state of the muscle. Needle muscle biopsies or phospho-nuclear magnetic spectroscopy can be used to study the accumulation and depletion of fatigue-related metabolites (i.e., decreasing intramuscular phosphocreatine concentrations [PCr] and increasing muscle lactate, adenosine diphosphate [ADP], inorganic phosphate [Pi] and hydrogen ion [H^+^] accumulation) during severe-intensity CWR tests (12, 13). However, while these methods provide clear insights into muscular metabolic changes, they are invasive, expensive, and not easily accessible, and are therefore traditionally impractical in the elite sporting environment. Alternatively, the temporal pulmonary rate of oxygen consumption (V^·^O_2_) profile may indirectly provide insights into the energetic state of the muscle (9, 13-15). For example, slower O_2_ onset kinetics can reflect a greater O_2_ deficit, which is associated with greater substrate-level phosphorylation and anaerobic metabolism (i.e., decreased [PCr] and increased blood lactate concentration), resulting in greater homeostatic disturbance and reduced exercise tolerance. Theoretically, assessing V^·^O_2_ kinetics during CWR tests can provide considerable insight into the energetic state of the muscle, allowing assessment of the effectiveness of training programs as a whole and other interventions (e.g., biomechanical).

Although the measurement of key physiological factors underlying test performances are essential, there is also a need to determine the most relevant outcomes relating to performance success: the functional performance outcome. Since the goal of a CWR test is to maintain a constant work output, fatigue assessment is problematic as no external power loss occurs (unless the limit of tolerance, or exhaustion, is reached). One solution is to perform a maximal, movement-specific sprint test immediately after a CWR test to assess the muscle’s capacity for maximal, explosive output (16-19). Greater decrements in maximal sprint power have been observed as the duration (30 s – 10 min) of prior exercise is increased (when performed at 60-98% of the mechanical power output at V^·^O_2max_) (16, 17, 20). Assuming that athletes are highly motivated to produce maximal effort and are familiar with the testing procedures, the functional loss in maximal sprint power immediately after a CWR test should provide information about the functional state of the muscles; and, therefore, information about the task fatigability.

Submaximal (60-90% of the mechanical power at V^·^O_2max_) exercise tests are commonly used to predict maximal cycling performance (e.g. YMCA, Astrand-Ryhming, Physical Work Capacity 170, etc.) (21, 22). Yet to truly translate controlled laboratory test results to field performance, the conditions under which the physiological limitations are manifested should be simulated (19). Maximal effort laboratory tests are therefore needed to accurately reflect competition intensities. However, highly fatiguing tests require hours, or days, of recovery due to resulting physiological and psychological deficits and may disrupt an athlete’s training schedule. Consequently, it is of interest to coaches and athletes to determine whether the data of shorter, less fatiguing tests provide comparable information to longer, more fatiguing tests.

The main purpose of this study was to determine whether the physiological variables measured during a shorter CWR test (2 min, 50% of the complete test duration) as well as the functional mechanical capacity measured immediately after it can be used as proxies for the fatigability results obtained in a longer (4-min) test. Because fatigue assessment is problematic during CWR tests, a secondary aim was to assess the relationship between the physiological changes, particularly V^·^O_2_ kinetics, measured during the CWR tests and the decline in maximal sprint power measured after the CWR tests. Cycling was chosen as the model within which to test the hypothesis as it is a widely used exercise testing and training modality with both sporting and clinical applications, as well as being task-specific to the track cycling population that formed the current study cohort. We hypothesised that both the measured physiological variables and functional decline in the shorter test would be of lesser magnitude, and be associated with the outcomes of the longer test.

## Materials and Methods

### Participants

Sixteen highly trained male (*n*=13) and female (*n*=3) cyclists (age 18.7 ± 2.2 y; height 180.8 ± 8.0 cm; mass 73.2 ± 10.1 kg; V^·^O_2peak_ 64.4 ± 6.0 ml·kg^-1^·min^-1^; estimated mechanical power at V^·^O_2peak_ 353.3 ± 58.9 W) volunteered to participate in the study. All participants were involved in an Australian State Institute of Sport Track Cycling program, completed ≥4 cycling and two resistance training sessions a week, and were free from injury at the time of data collection. All cyclists had ≥2 years’ experience in the high-performance environment and were familiar with high-intensity ergometer testing. Written informed consent was given prior to participation and the study was approved by the Human Research Ethical Committee of Edith Cowan University (project number 2019-00505-DUPLESIS).

### Data collection

#### Experimental design

Data were collected across two days separated by at least 48 h but at the same time of day using a stationary electromagnetically braked cycle ergometer (LODE Excalibur, Groningen, Netherlands). On Day 1, an industry-modified incremental step test (23) was completed including a submaximal component, a 20-min recovery, and a constant work rate (CWR) test of 4 min. This “2-in-1” test was designed to replicate a track cyclist’s pursuit time trial (i.e., an individual pursuit across 4 km for men, and 3 km for women). The purpose was to record the cyclist’s highest sustained average power output during the 4-min CWR test. The maximal rate of oxygen uptake (V^·^O_2peak_), V^·^O_2_ kinetics, heart rate, and blood lactate concentrations ([La^-^]_b_) were measured to indicate physiological stress, i.e., to infer the chemo-energetic state of the muscles across the 4-min CWR test. A 10-s maximal cycle sprint was performed before and immediately after the 4-min CWR test to assess the loss in maximal mechanical functional capacity (i.e., task fatigability). On Day 2, a shorter (2-min) CWR test was completed at the same intensity as the 4-min test, with a maximal 10-s sprint also performed before and after.

#### Experimental procedures

Day 1: “2-in-1” test

On Day 1, the seat, handlebar positions and crank length of the cycle ergometer were adjusted to match the participant’s own track racing bicycle. The participants were required to use their own pedals and cycling cleats as well as to perform a 15-min warm-up at a self-selected intensity, which was recorded and then repeated on Day 2. The warm-up procedures subsequently included a low-intensity bout at 50-100 W for 5 min, a 5-min ramp between 125-300 W, a 6-s maximal seated sprint, and finally a non-fatigued 10-s all-out seated sprint (PRE-CWR_,4min_), with each sprint separated by a 5-min active recovery at 50-100 W. All sprint components of the warm-up were completed in the cadence-dependent linear mode (i.e., in ‘Wingate’ isoinertial mode) of the ergometer with a torque factor load of 0.6 Nm·kg^-1^. All other warm-up components were completed using a constant power mode of the ergometer, which allows the power output to be set independent of the freely chosen cadence.

After the 15-min standardised warm up, participants performed a modified V^·^O_2max_ test, i.e., the “2-in-1” test, using methods currently employed in the State Institute system in Australia. This test comprised two components separated by 20 min of active or passive rest as selected by the participant (and repeated on Day 2). The first component involved 5-7 × 5-min stages of increasing intensity, starting at 100 W (women) or 150 W (men) and increasing by 25 W (women) and 50 W (men), using the constant power mode of the ergometer. Rating of perceived exertion (RPE (24)) was obtained at the end of each stage. Blood lactate samples were taken from the earlobe and analysed (Lactate Pro2, Kyoto, Japan) during the 4^th^ minute of each stage, and the test ceased when [La^-^]_b_ exceeded 4 mmol·l^-1^. V^·^O_2_ and carbon dioxide production (V^·^CO_2_) data were continuously collected using a ParvoMedics metabolic cart system (ParvoMedics TrueOne 2400 diagnostic system, USA) and averaged over 10-s windows during the last 2 min of each stage. Heart rate was continuously measured (RS800 Polar Heart Rate Monitor, Finland).

Following the 20-min recovery, a warm-up of 5 min at a self-selected intensity was mandated after which the second component of the test, a 4-min CWR test, was performed. This 4-min CWR test was designed to imitate a pursuit race in which bicycle gears are fixed and power output is highly cadence-dependent. Therefore, the CWR test was completed using the cadence-dependent mode of the ergometer with careful consideration of the torque factor load setting. Through consultation with the high-performance coaches and considering each participant’s recent (<6 months) race history, one of two processes was followed to determine test power output and cadence, and therefore the appropriate torque factor load:

For the minority of participants (*n*=5) who used power meters on their track bicycles, the target power and cadence were set to their most recent Individual Pursuit power output and cadence;

For the remaining participants, the V^·^O_2_ during the last 2-min of each submaximal workload of the “2-in-1” test was averaged. Based on the linear regression equation of the submaximal V^·^O_2_ and power output, the V^·^O_2peak_ (determined from the cyclists’ previous test conducted within the last year) was used to predict the mechanical power output at V^·^O_2peak_. The target power output of CWR test was estimated between 105-110% of this mechanical power output at V^·^O_2peak_, with the estimation based on previous research (25, 26) and retrospective analysis of all athlete data collected in the “2-in-1” test at the State Sports Institute in the three years preceding testing.

The torque factor load was subsequently calculated as:

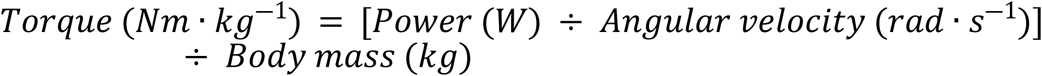

with power being the predicted 4-min CWR test power output, and angular velocity calculated as:

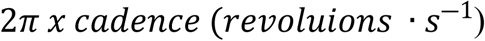

The participants were familiar with performing this severe-intensity CWR test and therefore strongly encouraged to provide a maximal but evenly-paced effort, and to maintain the target cadence for the duration of the test. The aim was to attain the highest average sustained power output, even if it resulted in a small cadence variation (of less than 5 rpm). A head unit secured to the ergometer provided visual feedback on both cadence and power. They were instructed to remain seated throughout the test. Torque and angular velocity (and power) data were sampled every 2° of a pedal stroke by the LODE ergometer mechanical system, and subsequently averaged per pedal revolution. The average power and cadence of the 4-min CWR test were calculated and used as reference for testing on Day 2. V^·^O_2_, V^·^CO_2_ and heart rate data were continuously collected throughout the 4-min CWR test.

At the immediate conclusion of the 4-min CWR test, without pause, a 10-s all-out seated sprint (a “fatigued sprint”, POST-CWR_,4min_) was performed (18, 19). No pause was allowed after the 4-min CWR test to prevent recovery of central or peripheral aspects of fatigue. A cycling-specific maximal sprint was selected to assess fatigue (i.e., rather than a knee extension, leg press or other test) to prevent any perseveration effect (i.e., a motor pattern interference) influencing the test performance, which may occur when movement patterns differ between tasks (27, 28). Blood was obtained from the ear lobe before and 1, 3, 5 and 7 min after the sprints that followed the CWR tests. [La^-^]_b_ was analysed within 15 s using the Lactate Pro2 analyser and the peak [La^-^]_b_ value retained for further analysis. RPE was obtained immediately following PRE-CWR and POST-CWR.

Day 2: 2-min CWR test

On Day 2, participants completed the pre-test preparations as on Day 1, including the same warm-up that ended with a 10-s all-out seated sprint (PRE-CWR_,2min_). After 5 min active recovery (50-100 W), participants were equipped with the same metabolic mouthpiece, and then performed a 2-min CWR test at the same intensity (i.e. the same torque factor, cadence and power) as the 4-min CWR test. This 2-min CWR test was also immediately followed by a 10-s all-out seated sprint effort (POST-CWR_,2min_). V^·^O_2_, V^·^CO_2_, heart rate, [La^-^]_b_ and RPE data were collected as in the 4-min CWR test.

### Data Analysis

#### Physiological variables

V^·^O_2peak_, minute ventilation (VE_peak_), the respiratory exchange ratio (RER_peak_), and heart rate (HR_peak_) were calculated as the highest average values attained over any 30-s interval during both the 2- and 4-min CWR tests. The mean response time (MRT) represented the time taken from the baseline to achieve 63% of the final response V^·^O_2_, and was estimated for both the 2- and 4-min CWR tests using a mono-exponential model fitted to each V^·^O_2_ data set, as follows (9, 15, 29, 30):

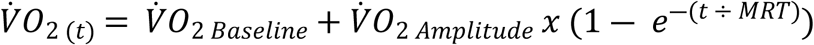

where V^·^O_2 (t)_ is the V^·^O_2_ at any time point, V^·^O_2 Baseline_ is the baseline V^·^O_2_ calculated as the 1-min average V^·^O_2_ before the start of the 2- and 4-min CWR tests, V^·^O_2 Amplitude_ is calculated as the difference between the steady state V^·^O_2_ asymptote and baseline, and (1-*e*^-(time ÷ MRT)^) is the exponential function that describes the rate of V^·^O_2_ rise towards the steady state amplitude. For each participant, the Microsoft Excel solver function was used as an iterative fitting procedure to solve for the smallest sum of squares differences between the projected V^·^O_2(t)_ (calculated using the exponential function) and the experimental data(9). V^·^O_2 Baseline_ was used as a fixed parameter and the V^·^O_2 Amplitude_ and the estimated MRT parameters for the 2- and 4 min CWR tests were subsequently computed.

The estimated MRT for the 2-min CWR test was factored since a steady-state (required for the exponential function above) could not always be clearly defined for V^·^O_2_ data collected during the 2-min CWR test. That is, in addition to calculating MRT for the 4-min CWR test from Day 1 (with a clearly defined steady-state), the first 2-min of the same V^·^O_2_ data set was used separately to calculate the 2-min MRT. The linear relationship (i.e., the slope and intercept) between the MRT of the first 2-min of the 4-min CWR test and the MRT of the 4-min CWR test was subsequently calculated for all individuals. This slope and intercept were then used to correct the MRT for the standalone 2-min CWR test from Day 2:

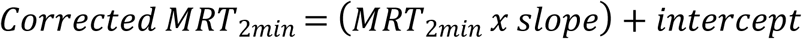

#### Mechanical variables

The functional decline in mechanical power output was determined by the fatigue-related decrements in peak and average power between the PRE-CWR and POST-CWR sprints. Only the first 8 s of the 10-s all-out sprints were used to remove any change in effort as the test neared completion. In addition, the peak cadence decrement during the sprints were calculated as indicators of the velocity-specific effect of the functional decline.

It should be noted that PRE-CWR was performed from a stationary start whereas POST-CWR was performed without delay from a rolling start following the 2- and 4-min CWR tests (i.e., the starting pedalling rates of the sprints were different). To eliminate the confounding effects of the pedalling rate on fatigue estimation, the average power was also calculated for a pedal rate-specific region (31). This region consisted of the minimal-to-maximal cadence range recorded during POST-CWR and compared to PRE-CWR within this region (see Fig 4).

### Statistical Analysis

All statistical analyses were performed using R software package (v 1.4). Data are presented as mean ± standard deviation, statistical significance was accepted at an alpha level of 0.05, and 95% confidence intervals (CI) with upper and lower limits were computed. Preliminary tests assessed and verified all test assumptions of multivariate normality (mvnormtest), multicolinearity (rstatix), homogeneity of variances (rstatix), and sphericity of the data, where relevant.

One-way repeated measures ANOVAs were used to test for differences between the 2- and 4-min CWR tests for i) the physiological variables obtained during the CWR tests [peak oxygen consumption (V^·^O_2peak_), estimated mean V^·^O_2_ response time (MRT), peak heart rate (HR_peak_), peak respiratory exchange ratio (RER_peak_), peak minute ventilation relative to body mass (VE_peak_), and peak blood lactate concentration ([La^-^]_b,peak_), as well as the perceived exertion (RPE)], and ii) changes in mechanical variables obtained during sprints performed before and after the CWR tests [peak power (ΔPower_peak_), average power (ΔPower_av_), average power calculated during the pedal-rate-specific region of POST-CWR (ΔPower_av,PRS_), and peak cadence (ΔCadence_peak_)].

Pearson’s correlations were computed to quantify the linear relationships between the physiological and mechanical variables between the 2- and 4-min CWR tests. These were interpreted as r: 0.10-0.39, weak; 0.40-0.69, moderate; 0.70-0.89, strong and 0.90-1.00 very strong relationship (32). The standard error of the estimate (SEE) of the regression was used to assess the accuracy of the 2-min test’s data to predict the outcomes of the 4-min test. Smaller SEEs represent a smaller prediction error. To assist in the interpretation of the statistical confidence, the relative SEE was presented as a percentage of the 4-min CWR test mean: SEE (%) = (SEE ÷ mean) × 100.

Relationships between the physiological and mechanical variables were analysed using a fixed-slopes linear mixed effect model approach using R package lmerTest (33). Visual inspection of residual plots confirmed that linear modelling assumptions were met. The response variable was the relative change in average power (W·kg^-1^) between the sprints across the 2- and 4-min CWR tests. The fixed effects were the physiological variables (V^·^O_2peak_, [La^-^]_b, peak_, MRT, HR_peak_, RER_peak_) as well as the duration (i.e., 2- and 4-min) of the CWR test. The random effect was set for the individual participants to account for intraindividual dependencies interindividual heterogeneity.

## Results

### 2-and 4-min CWR test outcomes

CWR tests were completed at 109 ± 0.1% of the predicted mechanical power output at V^·^O_2peak_ as determined from the incremental step test. Average power output and cadence for the 2- and 4-min CWR cycling tests were 384.0 ± 67.2 W (5.2 W·kg^-1^) and 106.6 ± 4.0 rpm vs. 383.4 ± 67.7 W (5.2 W·kg^-1^) and 105.5 ± 7.1 rpm, respectively.

### Physiological variables and perceived exertion between 2- and 4-min CWR tests

No differences were observed in V^·^O_2_ kinetics (including V^·^O_2peak_, and MRT) or RER_peak_ between the 2- and 4-min tests (p>0.05) (Table 1). Differences were observed in HR_peak_ (p=0.022), [La^-^]_b,peak_ (p=0.019), VE_peak_ (p=0.003) and RPE (p<0.001). Moderate to very strong correlations were found between 2- and 4-min CWR tests for all physiological variables (ranging from r=0.66 for MRT to r=0.96 for V^·^O_2peak_), but not for RPE (r=-0.21) (Fig 1). Relative to the 4-min CWR test mean values, SEEs ranged from 2.6% to 16.3%. Participants reached the same fraction of V^·^O_2peak_ at the end of the 2-min CWR test (93.4 ± 2.6%) as at the 2-min point of the 4-min CWR test (92.4 ± 2.8%) (Fig 2). In some cases, the 2-min CWR test was too short to clearly show a V^·^O_2_-time asymptote. Therefore, the MRT from the 2-min CWR test was corrected from 34.9 ± 6.1 s to 36.5 ± 5.1 s based on the linear relationship between the estimated MRT in the first 2-min of the 4-min CWR test and the estimated MRT of the 4-min test: [Corrected MRT_2min_ = 0.845 (MRT_2min_) + 7.02].

**Table 1.** Physiological variables and perceived exertion observed in the 2-and 4-min CWR tests (*n*=16).

|  | 2-min | 4-min | Mean Difference [95% CI] | p-value | r [95% CI] | SEE | SEE (%) |
| --- | --- | --- | --- | --- | --- | --- | --- |
| VO<sub>2peak</sub> (ml·kg<sup>-1</sup>·min<sup>-1</sup>) | 60.1 ± 5.9 | 64.4 ± 6.0 | 4.2 [3.3, 5.2] | 0.083 | 0.96 [0.88, 0.98] | 1.8 | 2.8 |
| MRT (s) | 36.5 ± 5.1 † | 35.5 ± 5.7 | -1.0 [-3.4, 1.3] | 0.767 | 0.66 [0.25, 0.87] | 4.4 | 12.4 |
| RER<sub>peak</sub> | 1.15 ± 0.07 | 1.16 ± 0.05 | 0.01 [-0.01, 0.03] | 0.500 | 0.77 [0.45, 0.92] | 0.03 | 2.6 |
| HR<sub>peak</sub> (bpm) | 181.5 ± 9.0 | 188.6 ± 8.5 | 7.1 [3.7, 10.5] | 0.022* | 0.75 [0.39, 0.91] | 5.8 | 3.1 |
| [La<sup>-</sup>]<sub>b,peak</sub> (mmol·l<sup>-1</sup>) | 9.5 ± 2.1 | 11.7 ± 2.9 | 2.2 [1.2, 3.1] | 0.019* | 0.76 [0.43, 0.91] | 1.9 | 16.3 |
| VE<sub>peak</sub> (l·kg<sup>-1</sup>·min<sup>-1</sup>) | 1.9 ± 0.3 | 2.3 ± 0.3 | 0.4 [0.3, 0.5] | 0.003* | 0.83 [0.57, 0.94] | 0.2 | 8.7 |
| RPE (6-20 Borg scale) | 15.8 ± 1.5 | 18.8 ± 0.7 | 3.0 [2.1, 4.0] | <0.001* | -0.21 [-0.64, 0.31] | 0.8 | 4.4 |

*Data presented as mean ± standard deviation. SEE: standard error of the estimate; smaller SEE represents a smaller prediction error. SEE (%): SEE relative to 4-min CWR* *test mean. MRT: Estimated mean response time; time taken to reach 63% of the VO<sub>2</sub> asymptote. †Corrected mean response time: MRT corrected by the relationship between* *the MRT of the 4-min CWR test and the first 2-min within the 4-min CWR test. \* $p<0.001$ , significant difference between sprints performed before and after the CWR tests.*

**Fig 1.**
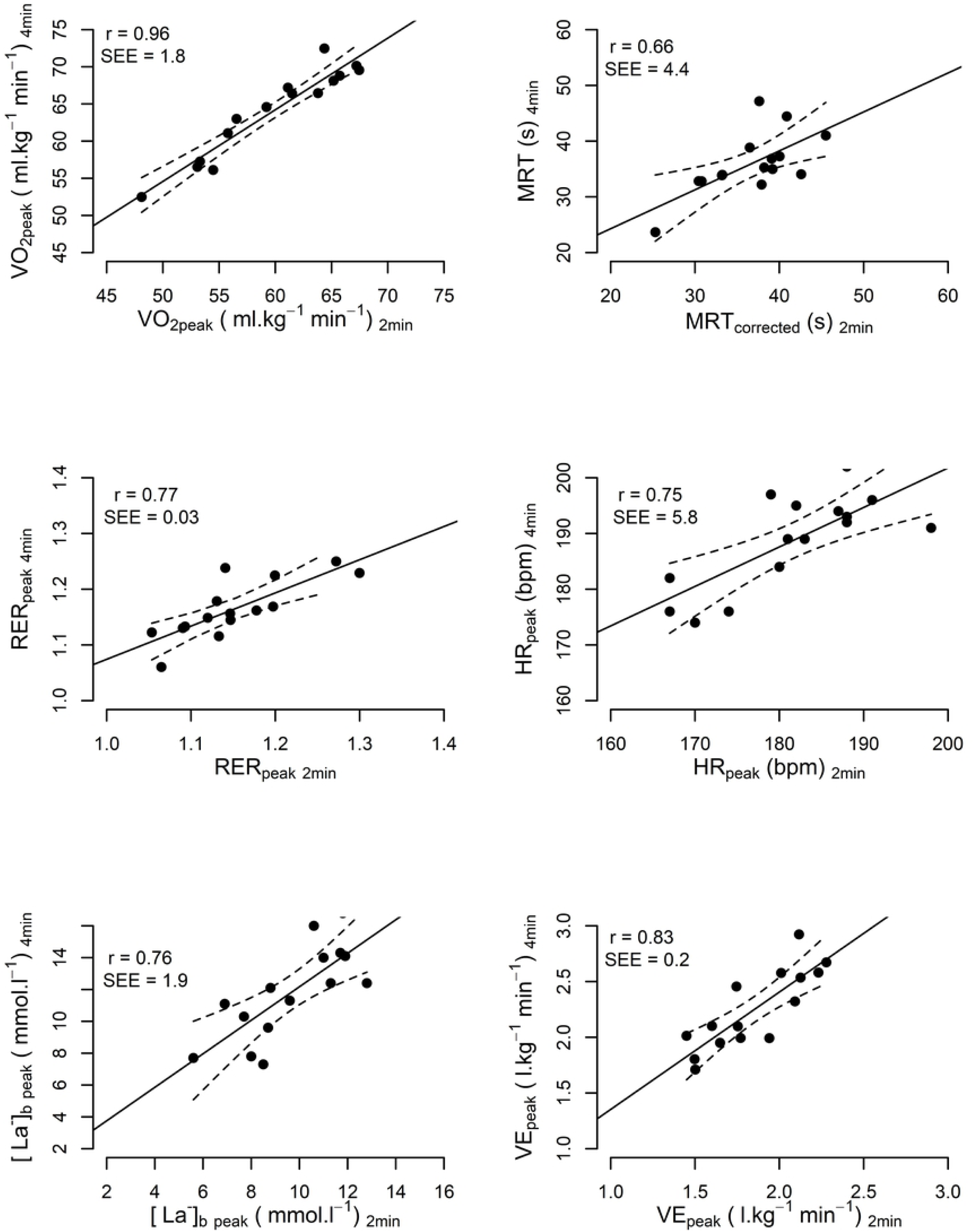
Relationships between physiological changes induced by 2- and 4-min CWR tests. Correlation and absolute standard error of the estimates (SEE) between the CWR tests are displayed on the top left corner of each graph. Dotted lines indicate the 95% CI. Physiological variables obtained during the 2-min CWR test show moderate to strong correlations and can predict the outcomes of the 4-min CWR test within a 2.6-16.3% error range (relative SEE), although a portion of this error relates to the tests being conducted on different days (see text for details).

**Fig 2.**
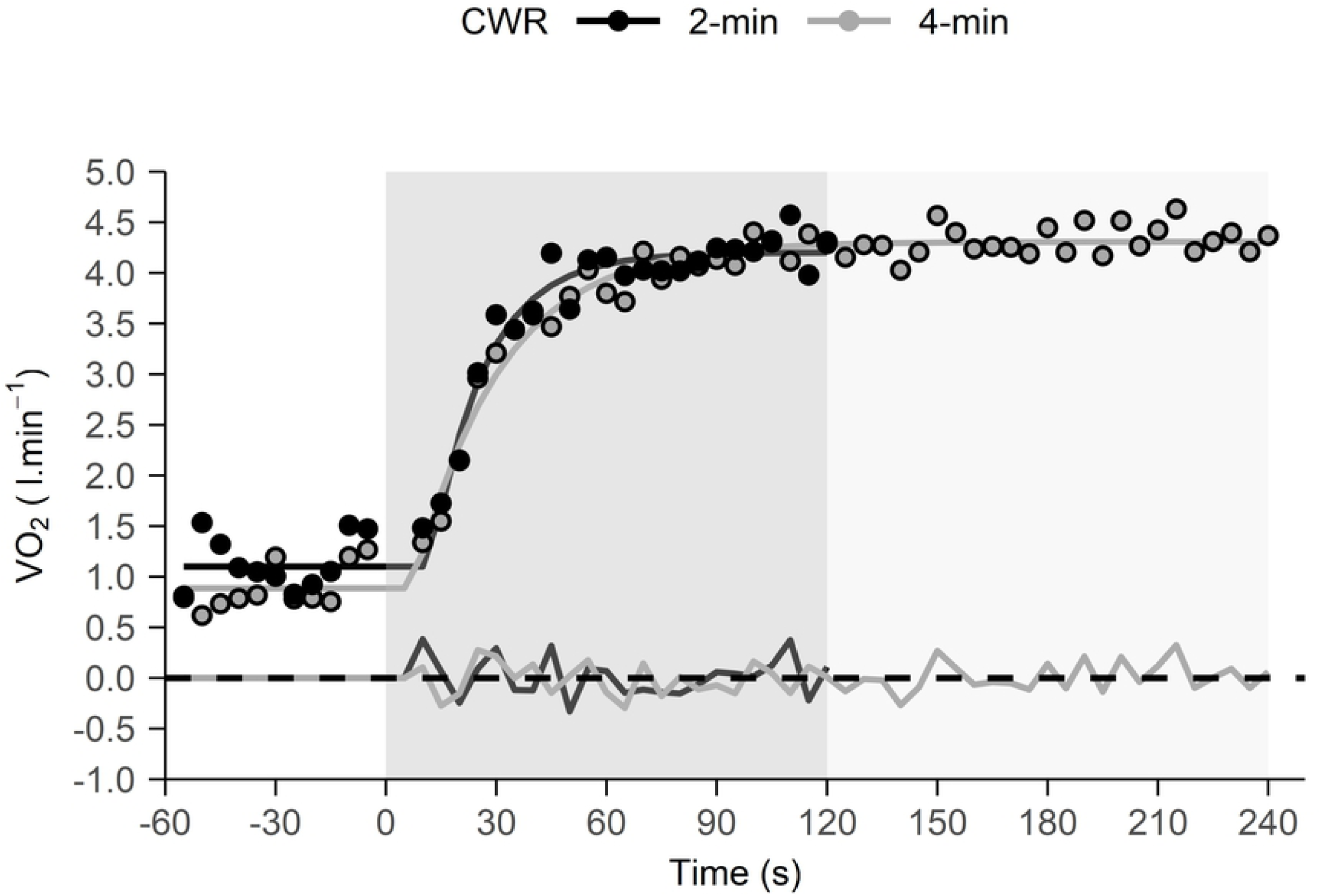
V^·^O_2_-time profiles for 2- and 4-min CWR tests in a representative participant performed on different days. V^·^O_2_ kinetics are inferred by the mean response times (Corrected MRT_2min_=25.3 s vs. MRT_4min_=23.6 s) and the V^·^O_2_ steady state, as fitted by a mono-exponential function which were similar for both tests. The bottom black and grey lines represent the residuals. The darker and lighter grey shaded regions show the 2- and 4-min CWR test durations, respectively.

### Mechanical variables of the sprints performed before and after the 2-and 4-min CWR tests

Changes (Δ) in mechanical variables from the sprints performed before and after the 2-min CWR test were significantly smaller (p<0.001) than changes in the sprints performed before and after the 4-min test (Table 2). Moderate to very strong correlations were observed in mechanical variables between the 2 and 4-min test (ranging from r=0.67 for ΔCadence_peak_ to r=0.92 for ΔPower_peak_) (Fig 3). Relative to the 4-min CWR test mean values, SEEs ranged from 14.2% to 26.1%. Fig 4 shows the power- and torque-cadence relationships of the sprints performed before and after the CWR tests and its downward shift in the fatigued conditions. Fig 5 shows the changes in peak power relative to body mass for the individual groups of cyclists. Irrespective of some individual variation, an overall trend of a functional mechanical decline in sprint capacity after the longer vs. the shorter test is indicated.

**Table 2.** Differences in the changes in mechanical variables recorded during sprints performed before (PRE-CWR) and after (POST-CWR) the 2-and 4-min CWR tests (*n*=16).

| | $\Delta$ Sprint<br>2-min CWR | $\Delta$ Sprint<br>4-min CWR | Mean Difference [95% CI] | p-value | r [95% CI] | SEE | SEE (%) |
| --- | --- | --- | --- | --- | --- | --- | --- |
| $\Delta Power_{peak}$ (W) | $-379.1 \pm 153.9^*$ | $-553.2 \pm 215.9^*$ | 174.1 [124.0, 224.3] | $< 0.001^{**}$ | 0.92 [0.79, 0.97] | 85.1 | 15.4 |
| $\Delta Power_{ave}$ (W) | $-305.2 \pm 83.2^*$ | $-455.8 \pm 126.1^*$ | 150.6 [111.1, 190.0] | $< 0.001^{**}$ | 0.83 [0.56, 0.94] | 73.5 | 16.1 |
| $\Delta Power_{ave,PRS}$ (W) | $-420.7 \pm 113.1^*$ | $-540.7 \pm 164.4^*$ | 120.0 [79.7, 160.2] | $< 0.001^{**}$ | 0.89 [0.71, 0.96] | 76.9 | 14.2 |
| $\Delta Cadence_{peak}$ (rpm) | $-28.6 \pm 9.1^*$ | $-49.0 \pm 16.7^*$ | 20.4 [13.7, 27.1] | $< 0.001^{**}$ | 0.67 [0.26, 0.88] | 12.8 | 26.1 |

Data presented as mean ± standard deviation. SEE: standard error of the estimate; smaller SEE represents a smaller prediction error. SEE (%): SEE relative to 4-min CWR test mean. $\Delta$ : changes between sprints performed before and after CWR tests. $Power_{ave,PRS}$ : average power during the pedal rate-specific region, i.e., specific to the cadence range of the fatigued sprint performed after the CWR tests. \*: significant difference between sprints performed before and after CWR tests, $p < 0.001$ . \*\*: significant difference between changes in sprints across the 2- and 4- min test durations, $p < 0.001$ .

**Fig 3.**
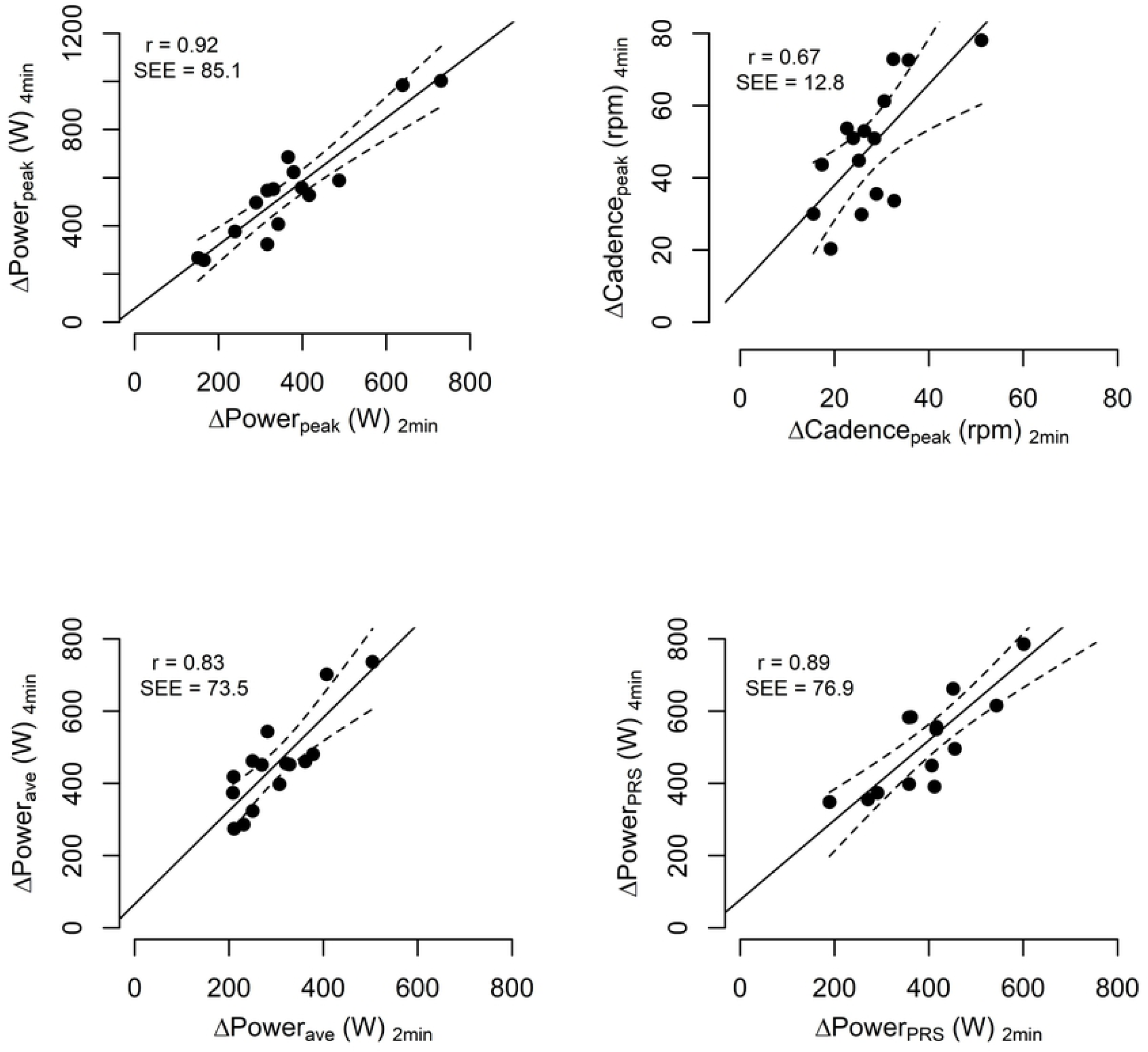
Relationships between 2-min and 4-min CWR tests for changes in (Δ) mechanical variables obtained in the sprints. Correlation and standard error of the estimates (SEE) between the CWR tests are displayed on the top left corner of each graph. Dotted lines indicate 95% CIs. Relative SEEs show that ΔPower (average, peak and within the pedal rate specific region) between the pre- and post-CWR sprints of the 2-min test can predict the outcomes of the 4-min test within 14.2-16.1% error, although a portion of this error relates to the tests being conducted on different days (see text for details).

**Fig 4.**
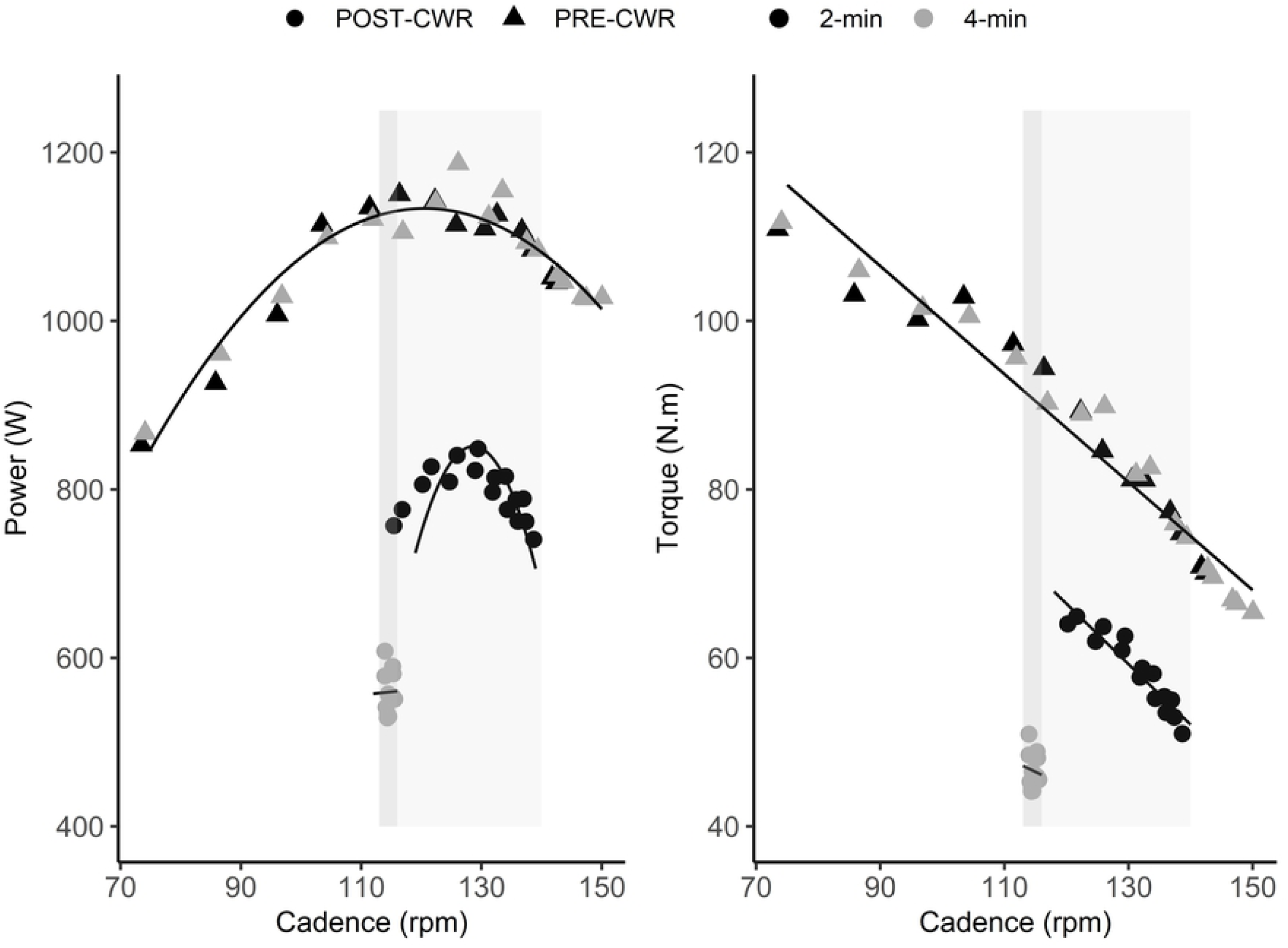
Power-cadence (left) and torque-cadence (right) relationships obtained in the sprints performed before and after the 2-min (black shapes) and 4-min (grey shapes) CWR tests from a representative subject. Triangles represent PRE-CWR sprints and the dots represent POST-CWR sprints. The downward shift of power- and torque-cadence relationships after CWRs demonstrate that power was severely compromised by the increase in CWR test duration. Shaded boxed regions represent the differences between the pedal rate ranges for POST-CWR_2min_ (lighter grey region) and POST-CWR_4min_ (darker grey region) zones used for ΔPower_ave, PRS_ calculation.

### Physiological variables of the CWR tests vs. changes in sprint mechanical variables

The linear mixed model was generated as: ΔPower_ave_ relative to body mass = (0.07 × V^·^O_2peak_) + (0.14 × [La^-^]_b,peak_) – (0.01 × MRT) + (1.65 × RER_peak_) + (0.02 × HR_peak_) – (1.26 × CWR test duration) -5.49, with a random effect for individual participants. An ANOVA of the model revealed significant effects for V^·^O_2peak_, [La^-^]_b,peak_ and CWR test duration (p<0.05). As a result, these physiological variables were found to be best related to the functional mechanical decline after the CWR tests.

## Discussion

The main purpose of this study was to determine whether a shorter (2-min), severe-intensity constant work rate (CWR) cycling test could be used as a proxy for a longer (4-min) test. This CWR test was selected to simulate a 4-km pursuit time trial in track cycling where performance is influenced by the ability to mitigate reductions in the power output and tolerate the inevitable peripheral muscle fatigue developed at these high physiological work rates. The findings confirmed our hypotheses. First, the CWR test of 50% of the original duration was sufficient to elicit substantial fatigue, as indicated by the significant loss in mechanical power, and this magnitude of reduction was strongly correlated with the power loss measured in the longer test. Additionally, both the physiological and mechanical variables (i.e., the fatigability estimates) were independently and strongly correlated between the CWR tests despite the tests being performed on different days, and thus being influenced by between-day variability. This highlights the comparable nature of the tests. Second, the differences in V^·^O_2peak_, blood lactate concentration, and the duration of the CWR tests were strongly associated with the decline in mechanical power output measured across the sprints. Although no single physiological variable can be used to predict the loss in mechanical power (i.e., fatigability) independently, explosive sprints both before and after a shorter CWR test can allow an estimate of fatigue that would be obtained in the longer test, without causing the same extent of fatigue. Shorter CWR tests followed by maximal sprints may therefore be useful as a (regular) fatigue assessment and monitoring tool in athletic testing environments.

The participants completed 2- and 4-min CWR tests at 109% of their predicted mechanical power output at V^·^O_2peak_, which reflected their individual track cycling pursuit power capacity. A decline of 32 ± 8% and 47 ± 9% in mechanical power (i.e., functional capacity) was caused by the 2- and 4-min CWR tests, respectively. Analogous to our findings, others (16, 17, 20, 34-36) have shown maximal power reductions of 25-32% after submaximal CWR cycling tests performed at 60-98% of the mechanical power reached at V^·^O_2peak_ for durations of 3-10 min. It is likely that the greater (47%) decrease in power output after the 4-min CWR test in the present study reflects the higher workloads (i.e., 109% vs. 60-98% of the mechanical power output at V^·^O_2peak_) and cadences (100-110 rpm vs. ∼60-90 rpm), higher level of athletic training ability, and potentially, the different sprint modality (i.e., isoinertial vs. isokinetic, as discussed below) used in the present study.

In addition to changes in functional mechanical power output, differences in the torque-angular velocity (or cadence) and power-angular velocity relationships during maximal cycling present valid estimates of performance fatigue (31, 36, 37). As done by Capelli and colleagues (17) and Marcora and Staiano (8), we employed a cadence-dependent mode (i.e. an isoinertial mode) on the cycle ergometer, allowing the participants to accelerate to a maximal cadence in each sprint. As illustrated in Fig 4, the linear torque-cadence and parabolic power-cadence relationships of the fatigued sprints shifted downwards, indicating that the athletes’ functional performance abilities were severely compromised by the prior CWR tests; i.e. they were highly fatigued (8, 31, 37). This was particularly evident after the 4-min CWR test. Evidently the participants were unable to re-generate the same level of torque and angular velocity after the CWR tests despite producing maximal voluntary effort. One may therefore gain insight into task fatigability by imposing a maximal sprint immediately after a CWR test, which highlights that greater performance fatigability was induced by the longer CWR test. This fatigability would otherwise not have been quantified since there was no mechanical drop in power during CWR tests. More importantly, the shorter test was discernibly less strenuous and should thus have less impact on subsequent recovery and training. This possibility should be explicitly determined in a future study.

As the cycle sprints were used to predict the functional loss rather than muscle fatigue specifically, sprint cadence was not fixed to the cadence of the CWR test. Thus, the velocity-dependent effect of fatigue was not accounted for (34, 38). Because a cadence-dependent mode was used, participants could increase cadence as a means of increasing power, which reflects the temporal and kinetic patterns obtained when using fixed gears on a track bicycle (where cadence must increase when power increases). Not only did the CWR tests compromise the maximal cadence achieved compared to a non-fatigued sprint, but the acceleration was also severely impacted. This is evidenced by similarities between the average CWR test cadence (105.5 ± 7.1 rpm) and peak sprint cadence (105.5 ± 11.1 rpm), which may partly be explained by the non-fatigued sprint commencing from a stationary start whilst the fatigued sprint commenced, without pause, at the final cadence of the CWR test (i.e., from a rolling start). To account for this limitation, we calculated the average difference in mechanical power for the ‘pedal rate-specific region’: i.e., the cadence range of the fatigued sprint. However, like the decline in peak power (32% and 47%), decrements in average power during the pedal rate-specific region were found after the 2-(37 ± 5%) and 4-min (51 ± 6%) CWR tests. Therefore, the functional performance outcomes were severely affected by both 2-and 4 min CWR tests even when partly accounting for velocity-dependent differences.

As illustrated by the loss in functional mechanical power, the present results show that a CWR test of 50% of the original duration was sufficient to induce fatigue, regardless of the exact mechanisms underpinning the fatigue (i.e., in the muscular system, cardiovascular system, or central nervous system) (11). Moreover, strong correlations (r=0.83-0.92) were found between the functional decline in power induced by the 2 and 4-min CWR tests within a prediction error (SEE) of 14-16%. (Fig 3). Considering that the tests were done on the separate days, part of the SEE can be attributed to *between-subject* differences in the rate of fatigability (Fig 5) but the remainder can be attributed to the additional *between-day* variability of the sprint tests. For example, the PRE-CWR sprint alone showed a SEE of 2.7% (26.3 W difference in average power, r=0.99) between days whereas the fatigued POST-CWR showed a higher SEE of 8.5% (42.6 W difference in average power, r=0.91). Therefore, the 16% prediction error of the change score (i.e., the fatigue estimate) will be elevated due to the between-day variability in the sprints themselves. It is also important to consider that tests done in the fatigued state also show greater variability. Previous researchers (39, 40) showed good reliability (SEE <5%, r >0.9) both within and between days for peak and mean power achieved in repeated maximal explosive ergometer tests, however their estimated fatigue indices indicated poor reliability (r=0.43 or ICC= 0.34 and CV= 21.3%). Due to the strong association between the 2-and 4-min CWR tests obtained in the present study, despite the tests being done on different days, one can be confident that the estimated functional decline estimated in the 2-min test is strongly reflective of that which would be obtained in a 4-min CWR test. Further research is required to specifically assess the systematic error between tests when they are performed on the same day with sufficient recovery, which was not possible in the present study.

**Fig 5.**
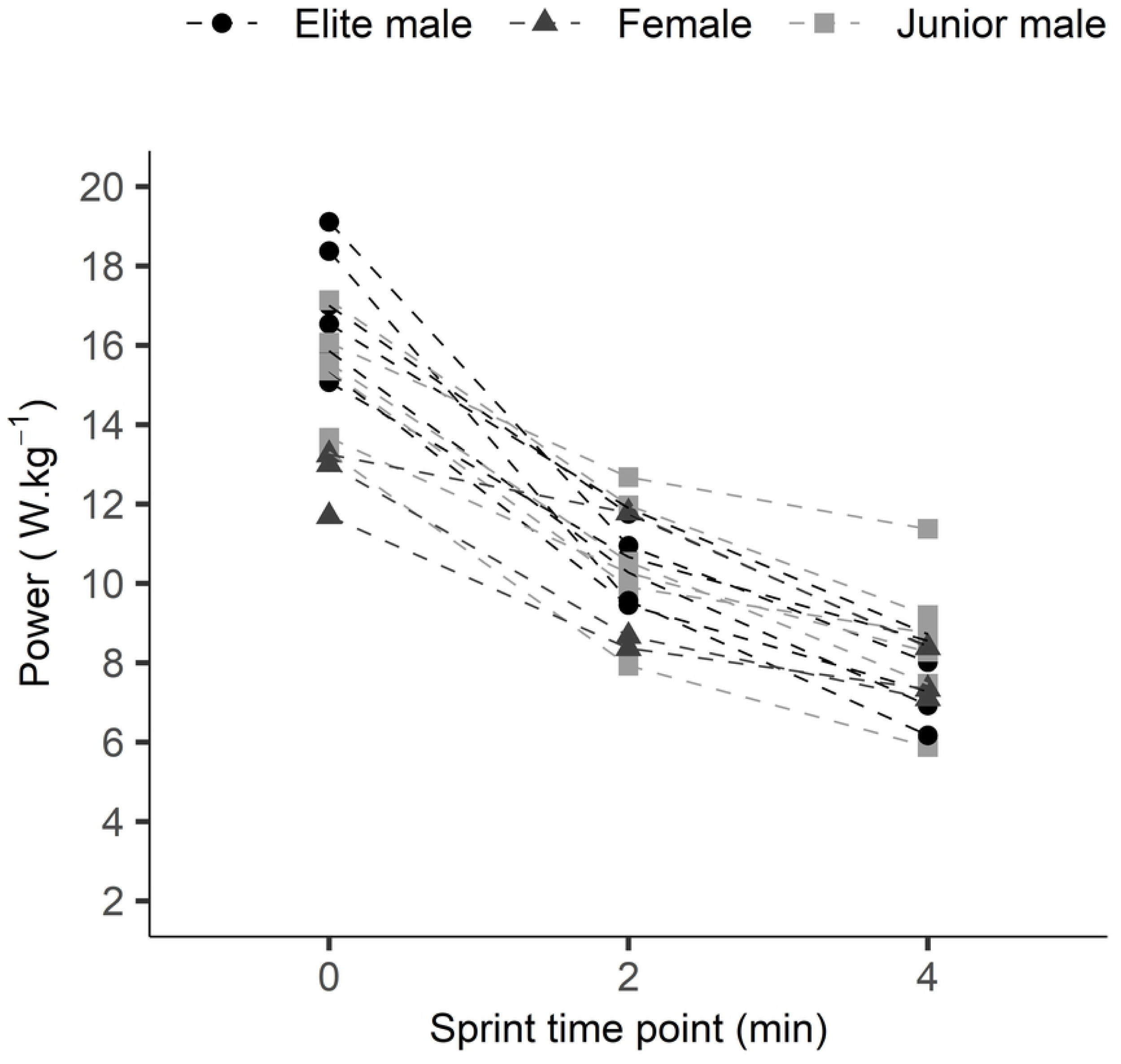
Peak power (relative to body mass) before and after the 2- and 4-min CWR tests for all participants. Irrespective of individual variation, a greater overall functional deficit was observed for the longer test duration. Sprint time point ‘0’ represents the relative peak power in pre-CWR sprints whereas time points ‘2’ and ‘4’ represent the relative peak power in sprints performed after the 2-min and 4-min CWR tests, respectively. Shapes represent the different groups of cyclists: black dots = elite men, light grey squares = junior (U19) men, dark grey triangles = women.

In addition to the power loss, an important aim was to assess differences in the physiological demands between the CWR tests to provide insight into the energetic state of the muscles and their influence on the subsequent maximal mechanical power loss. All physiological variables increased in magnitude with a similar trajectory between the tests, which eventually resulted in a greater decrease in mechanical power after the 4-min CWR test. Other researchers have found that exhaustive cycling exercise within the severe-intensity domain, regardless of work rate or test duration, is associated with the same level of depletion of high energy phosphates [PCr] and the accumulation of [H^+^], [ADP] and [Pi], as well as lactate due to the greater rate of glycolysis, and that the V^·^O_2_ will continue to rise until reaching V^·^O_2max_ (i.e. V^·^O_2_ slow component) (4, 10, 12, 13). This resulting muscular homeostatic imbalance can influence various components within the sarcomeric excitation-contraction coupling process, resulting in a functional decline manifested as a reduced external power output. As these methods are impractical for use within the elite sporting environment, alternative methods such as the temporal pulmonary V^·^O_2_ profile may provide insight into the energetic state of the muscle during severe-intensity exercise.

V^·^O_2_ kinetics (i.e., V^·^O_2peak_ and estimated mean response time), peak heart rate and blood lactate concentrations were measured as physiological stress indicators, i.e., to infer the chemo-energetic state of the muscles across the 2- and 4-min CWR tests. Strong relationships and small SEEs indicated that the physiological variables obtained in the 2-min CWR test can be used to predict the outcomes of the 4-min CWR test, at least within a 2.6-16.3% error range (Fig 1). It was identified that neither the duration of the CWR test nor the fact that the CWR tests were performed on different days affected V^·^O_2_ kinetics (and therefore the oxygen deficit) or RER_peak_. Alternatively, significant increases were found in peak blood lactate concentration, peak heart rate and minute ventilation as well as RPE for the longer CWR test. It can be assumed that the greater rise in these physiological variables during the 4-min CWR test would be associated with a greater anaerobic metabolism (i.e., decreased [PCr] and increased muscle lactate concentrations and therefore [H^+^] accumulation). This would result in a greater homeostatic disturbance and reduced exercise tolerance, or in our case lead to a greater change in mechanical power output measured immediately after the 4-min CWR test. Consequently, for the same power output within the severe intensity domain, exercising for a longer duration resulted in a greater instability of the internal metabolic environment, which was subsequently expressed as a greater loss in functional explosive power.

The findings from the mixed linear regression model suggest that V^·^O_2peak_, peak blood lactate concentrations and the duration of the CWR tests were significantly associated with the decline in the average power measured across the sprints. Thus, irrespective of the physiological variables measured in the 4-min test being strongly predicted by the outcomes of the 2-min test, no single physiological variable could predict mechanical power loss (i.e., fatigability) independently. Alternatively, pre- to post-CWR changes in explosive sprint performance may provide a useful fatigue estimate for use in training monitoring in athletic environments. Furthermore, whilst the change in sprint power from before to immediately after a shorter CWR test can be used to predict (at least within a 14-16% error range) the power loss induced by the longer CWR test, the opportunity also exists to use the post-CWR power alone (with SEE of 8.5%) as long as one is confident that the athlete started the CWR in the same state as the previous test; variability of a complex variable (e.g. post – pre score) should always be greater than a simple variable (e.g. post score alone).

Whilst taking into account the predictive error, a shorter CWR test may provide a valid substitute for the longer tests, which are significantly physiologically and psychologically taxing and induce substantial residual fatigue. Although the exact temporal recovery response remains to be tested in future studies, the shorter test was more tolerable as it induced less fatigue, and is not expected to substantively impact long-term athlete fatigue or training schedules. Consequently, it may be more frequently used in a training season (including at the start of a training session) to monitor performance and fatigue levels. The shorter test also introduces the possibility of completing multiple shorter tests within a single session in order to assess the effects of various interventions (such as bicycle-set up variations, recovery or nutritional interventions, etc.) and thus with reduced error imposed by between-day changes.

In conclusion, severe-intensity CWR tests of fixed duration may simulate track cycling individual pursuit performances, however a CWR test in itself rarely allows estimation of performance fatigue (unless the test results in task failure). Alternatively, fatigability can be estimated as the change in maximal sprint power from before to after a CWR test. The present results demonstrate that the physiological variables measured during, and the loss in mechanical (cycling) power measured immediately after, a severe-intensity 2-min CWR test may be used as a proxy for outcomes that would be obtained in a test of twice the duration (4 min) despite the test evoking less fatigue. Because the 2-min test is significantly less fatiguing, and thus more physiologically and psychologically tolerable, recovery time should be reduced and training and competition preparations less impacted; the specific magnitudes of these effects require further study. A shorter test might therefore be employed in athlete (or clinical) environments to minimise the ongoing impact of testing, and may speculatively allow for multiple tests to be completed within a single session, assuming adequate recovery time is provided, to assess the acute effects of specific interventions without between-day effects on test reliability.

## Conclusion

The CWR test lasting 50% of the original duration (i.e. 2 min vs. 4 min) was sufficient to evoke fatigue by detecting meaningful changes in maximal cycling power output, and both physiological and psychological strain without overly taxing the athletes. Laboratory tests that simulate competition intensities (such as an individual pursuit) can therefore be performed more regularly by using a shorter-duration CWR test that subsequently requires less recovery time and thus impacts subsequent training and competition preparation to a lesser extent. Such tests could be used to monitor training adaptation or to assess the effects of acute and chronic interventions, such as changing bicycle-rider biomechanics, recovery or nutritional strategies, equipment, etc. In well trained athletes, it is likely that multiple tests could be conducted on the same day, assuming adequate recovery is provided, increasing testing efficiency. Additionally, as the capacity to produce and maintain high power outputs after a prior fatiguing exercise bout is essential for performance success, the ability to produce higher power output for a given level of fatigue or produce the same power after a CWR of higher mean power would indicate a performance improvement. Importantly, end-burst power following sustained, high-power cycling has implications for other track (e.g. Keirin or bunch races) and road cycling race events where a final sprint often dictates race outcomes.

## Statements and Declarations

The authors have no conflict of interest or funding to declare in the publication of this manuscript.

## Acknowledgement

The authors would like to thank all participants for volunteering their time to complete the study. We also thank Kirstin Morris and Katie McGibbon for valuable feedback on a draft version of the manuscript, and Shih Ching Fu for his statistical advice. This work was supported by the Industry Engagement Scholarship between the Queensland Academy of Sport’s Sport Performance Innovation and Knowledge Excellence unit and Edith Cowan University.

## Notes

### Competing Interest Statement

The authors have declared no competing interest.

